# Selective and Controllable Trapping of Single Proteins in Nanopores using Reversible Covalent Bonds

**DOI:** 10.1101/2025.09.11.675362

**Authors:** Yuanjie Li, Saurabh Awasthi, Peng Liu, Anna D. Protopopova, Michael Mayer

## Abstract

Analysis of individual proteins using nanopores makes it possible to determine their size and shape in a label-free approach, within minutes, and from μL sample volumes. Short residence times of proteins in the nanopore, high electrical current noise, and bandwidth limitations of the recording electronics during resistive pulse recordings, however, limit the accuracy of size and shape analysis of individual proteins. The work presented here introduces a polymer surface coating of solid-state nanopores to minimize non-specific interactions of proteins with the nanopore wall while functionalizing it covalently with phenylboronic acid (PBA) groups. These PBA groups make it possible to trap selectively glycated proteins by taking advantage of the formation of reversible covalent bonds between PBA and vicinal diol groups of glycated amino acid residues on proteins. Dwell time analysis revealed two populations of resistive pulses: short pulses with dwell times *t*_*d*_ below 0.4 ms from free translocation of proteins and resistive pulses that we term “long events” that last from 0.4 ms to 2 s and result from intended transient covalent bonds between glycated proteins and PBA groups in the nanopore lumen. The choice of applied potential differences during nanopore recordings or the pH value of the recording buffer makes it possible to control and extend the most probable trapping time of proteins in the nanopore within one to two orders of magnitude. This approach provides the highest accuracy for the determination of protein volume and shape achieved to date with solid-state nanopores and reveals that a trapping time of 1 to 20 ms is ideal to achieve reliable volume and shape analysis while retaining high throughput of the analysis. This approach, hence, extends the residence time of natively glycated proteins or of proteins that are intentionally glycated by straightforward incubation in a glucose solution, thereby providing selectivity and improving the accuracy of nanopore-based characterization of single proteins.

## Introduction

Nanopores have emerged as powerful tools for label-free single-molecule analysis of DNA^1, 2^, RNA^3, 4^, peptides^5–8^, and proteins^9^. In particular, solid-state nanopores with diameters of 10s of nanometers enable the label-free detection of intact proteins, including folded proteins, protein complexes, and protein aggregates.^10–20^ For instance, measuring changes in ionic current as individual proteins move through an electrolyte-filled nanopore makes it possible to characterize their volume and shape.^10, 12, 21, 22^ Despite significant progress, however, challenges remain in optimizing nanopore-based protein characterization.

The first major challenge is the tendency of proteins to interact non-specifically with the walls of solid-state nanopores, leading to pore clogging and inaccuracy in the determination of protein shape.^21, 23–25^ Estimation of protein shape requires that the protein be sampled in as many different orientations as possible during its passage through the nanopore.^12^ To address this issue, various surface modifications have been explored for rendering the lumen of solid-state nanopores inert towards protein adsorption.^26–31^ For example, previous work by our group^21^ and others^32^ demonstrated the effectiveness of polymer coatings such as poly(acrylamide)-g-poly(ethylene glycol) (PAcrAm-g-PEG) and poly(acrylamide)-g-poly(ethylene glycol)-poly(2-methyloxazoline) (PAcrAm-g-PEG-PMOXA) for reducing protein adsorption.^23^

The second major limitation is the short residence time of proteins within the nanopore.^33^ Typically, the most probable dwell times for the free translocation of proteins through solid-state nanopores are below 10 μs, and only a small fraction of resistive pulses extend to 150 – 400 μs.^12, 25, 29, 34, 35^ Due to bandwidth limitations of the recording equipment, the amplitudes of short resistive pulses are often not completely resolved, constraining the signal-to-noise ratio and compromising precise characterization of protein volume and shape.^12^ Such short residence times also preclude the possibility of monitoring conformational changes of single proteins while they reside in the pore. Several strategies have been developed to prolong the residence time of proteins inside nanopores.^10, 33, 36–39^ In a recent elegant study, Schmid et al.^33^ utilized a porous DNA origami sphere to create an electro-osmotic trap by physically blocking the nanopore exit. This approach achieves single-protein trapping for hours, with the advantage that protein modification is not required. Possible limitations of this approach are that protein trapping is indiscriminate, and the positioning of the DNA origami particles can increase the baseline current noise.^19^ Another approach by Wei et al. employed chemical functionalization of the nanopore lumen by nitrilotriacetic acid (NTA) moieties for specific binding of His-tagged proteins.^36^ To succeed, three NTA groups next to each other were required to achieve sufficiently long trapping times in the high ionic strength environment of nanopore recording buffers that are typically used for resistive pulse recording. Closest to the approach presented here is work by Tang et al,^40^ who reported the use of 4-mercaptophenylboronic acid groups on gold-coated nanopipettes for selective sensing of glycoproteins. This approach was applied for label-free detection of IgG and demonstrated the potential for detecting glycoproteins in the presence of non-glycated proteins. Boronic acids and their derivatives are well-established molecular sensors due to their selective, reversible reaction with vicinal diols, a functional group commonly found in sugars, polysaccharides, as well as glycated and glycosylated proteins.^41–43^ Recently, PBA groups have been employed in biological nanopores to discriminate between different sugar molecules.^44–46^

Here, building on the benefits of polymer coatings to minimize non-specific protein adsorption in solid-state nanopores,^21^ we introduce the use of azide functional groups in the polymer (PAcrAm-g-PEG-Azide) and functionalize them in a one-step reaction with DBCO-activated phenylboronic acid (PBA) groups using strain-induced click chemistry.

By analyzing resistive pulses from single proteins in the PBA-coated solid-state nanopores, we demonstrate that glycated proteins could be trapped selectively with median dwell times that can exceed those of their non-glycated counterparts by two orders of magnitude. The resulting prolonged trapping times improved the accuracy of volume and shape estimations of single proteins to the best values that could thus far be achieved with solid-state nanopores. Moreover, this work reveals for the first time experimentally the required minimum residence time of single proteins in a nanopore to achieve close to maximal accuracy of estimates of their size and shape.

## Results and Discussion

### Functionalization of nanopores with PBA

We developed a straightforward and robust two-step “dip and rinse” approach to functionalize nanopores in SiN_x_ membranes with PBA (**Figure 1A**). In the first step, we incubated the nanopore chip in a 1 mM solution of HEPES buffer with a pH of 7.4 containing 0.1 mg/mL PAcrAm-g-PEG-Azide polymer for 1 h. The second step was to thoroughly rinse the resulting azide-coated nanopore chip with ultrapure water and then incubate it in a solution containing 160 mM NaCl, 10 mM HEPES, pH 7.4, and 1.2 mg/mL DBCO-PEG_4_-PBA for 1 h, followed by a thorough rinse with ultrapure water to remove unbound molecules. We confirmed the efficiency of the click reaction between azide groups on the polymer and DBCO-PEG-PBA in solution using UV-vis spectroscopy (**Supplementary Figure S1)**.

**Figure 1.**
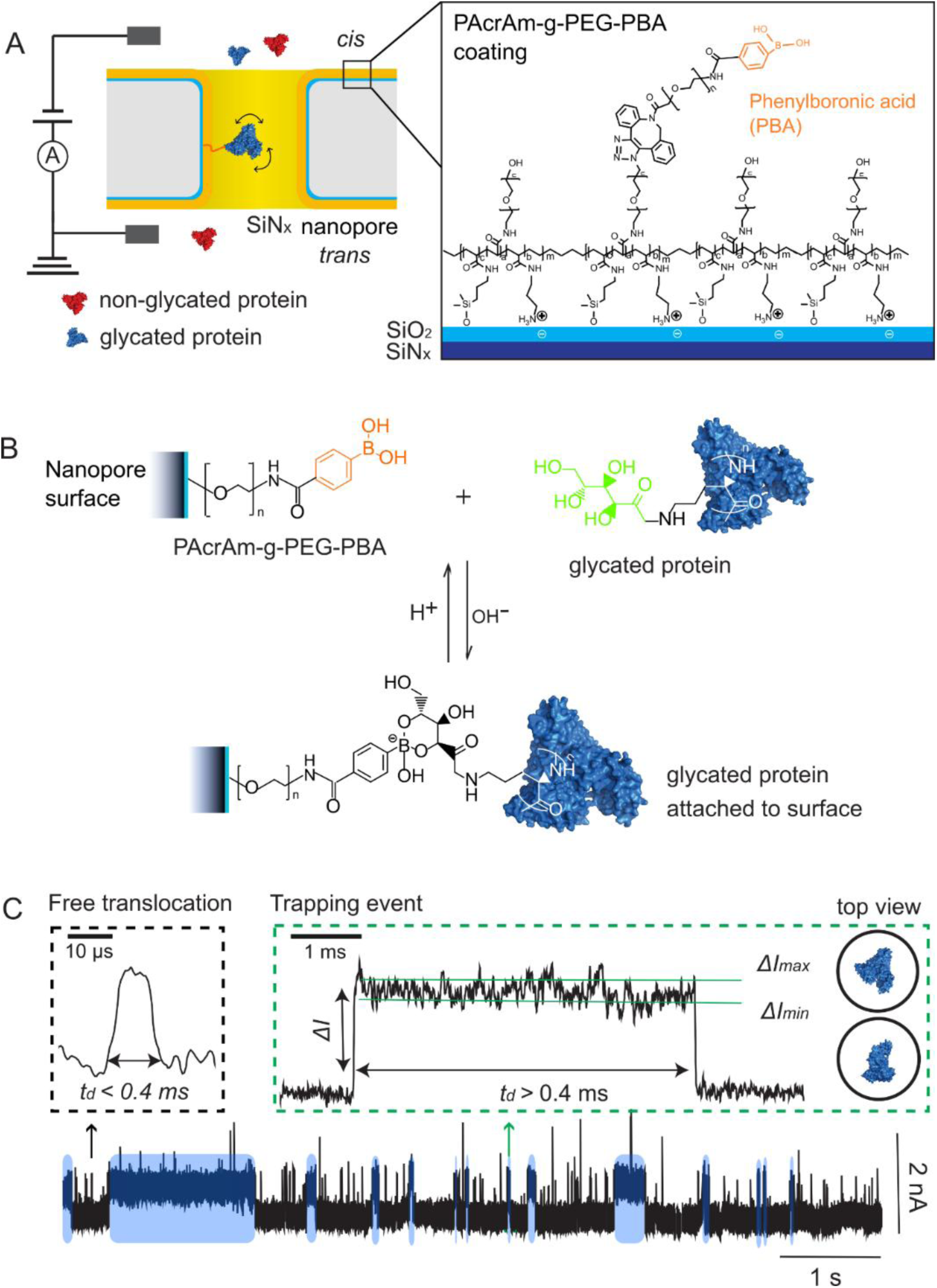
Reversible covalent bonds between glycated proteins and phenylboronic acid (PBA)-coated nanopores increase the duration of resistive pulses. **A)** Schematic of the polymer coating made from PAcrAm-g-PEG-PBA on the wall of a single nanopore in a SiN_x_ membrane. PBA groups transiently react with and trap glycated proteins (blue) while non-glycated proteins (red) translocate through the nanopores freely. **B)** PBA groups selectively capture glycated proteins via a pH-dependent reversible reaction with vicinal diols on carbohydrates attached to proteins. **C)** Representative ionic current trace in the presence of glycated human serum albumin (gHSA) in a PBA-coated nanopore with a diameter of 20 nm. Short upward spikes correspond to short resistive pulses (black inset), and prolonged current steps correspond to the formation of reversible covalent bonds (green inset). Some of the long resistive pulses induced by reversible covalent bonding (*t_d_* ≥ 0.4 ms) are indicated with blue shading. The *ΔI_min_* and *ΔI_max_* values of these long resistive pulses reflect the two extreme orientations of gHSA inside the nanopore: fully crosswise and fully lengthwise with respect to the long axis of the pore, as illustrated by the top-view cartoons of gHSA in the nanopore.^1^

We used ellipsometry and ion conductance measurements to estimate the thickness of the resulting PAcrAm-g-PEG-PBA coating. Ellipsometry results indicated that the average thickness of the dried PBA coating on the SiN_x_ chips was 1.1 ± 0.2 nm (*N* = 3, **Supplementary Figure S2A**). Ion conductance measurements across chips with nanopores of different diameters estimated an average thickness of the hydrated PBA coating of 1.4 ± 1.3 nm (*N* = 50, **Supplementary Figure S2B, Supplementary Note S1**).

In aqueous solution, PBA groups react with molecules containing vicinal diols, such as carbohydrates, by the formation of a boron-diol complex between phenylboronic acid and vicinal diols (**Figure 1B**).^42^ This reaction is reversible via the exchange of protons (H^+^) and hydroxide ions (OH^−^). To demonstrate the carbohydrate-binding activity of the PBA coating inside a nanopore, we measured the ion current through the nanopore in the presence of increasing concentrations of glucose in the recording buffer. The open pore current decreased as glucose molecules bound to the nanopore lumen and eventually approached a final value that corresponds to an estimated effective average thickness of the additional glucose coating of 0.83 ± 0.05 nm (*N* = 3) (**Supplementary Figure S3, Supplementary Note S1, S2**).

To test protein trapping with PAcrAm-g-PEG-PBA-coated nanopores, we translocated glycated human serum albumin (gHSA) or its non-glycated version HSA as a control through a PBA-coated nanopore with a diameter of 20 nm (**Figure 1C**). **Figure 2A** shows that the dwell time of most resistive pulses was significantly shorter than 400 μs, we attribute these pulses to free translocation of glycated HSA molecules that did not react with the PBA groups in the pore.^12^ **Figure 1C** and **Figure 2A,B**, however, also demonstrate that experiments with PBA-coated nanopores and glycated HSA displayed unusually long resistive pulses (*t_d_* > 400 μs) in addition to the short resistive pluses from free translocation. We hypothesized that these prolonged events result from the formation and dissociation of reversible covalent bonds between the glycated protein and PBA; these long pulses were absent in experiments with non-glycated HSA (red data in **Figure 2A, B**) as well as in experiments with nanopore coatings that lacked PBA groups (**Figure 2C, D**).

**Figure 2.**
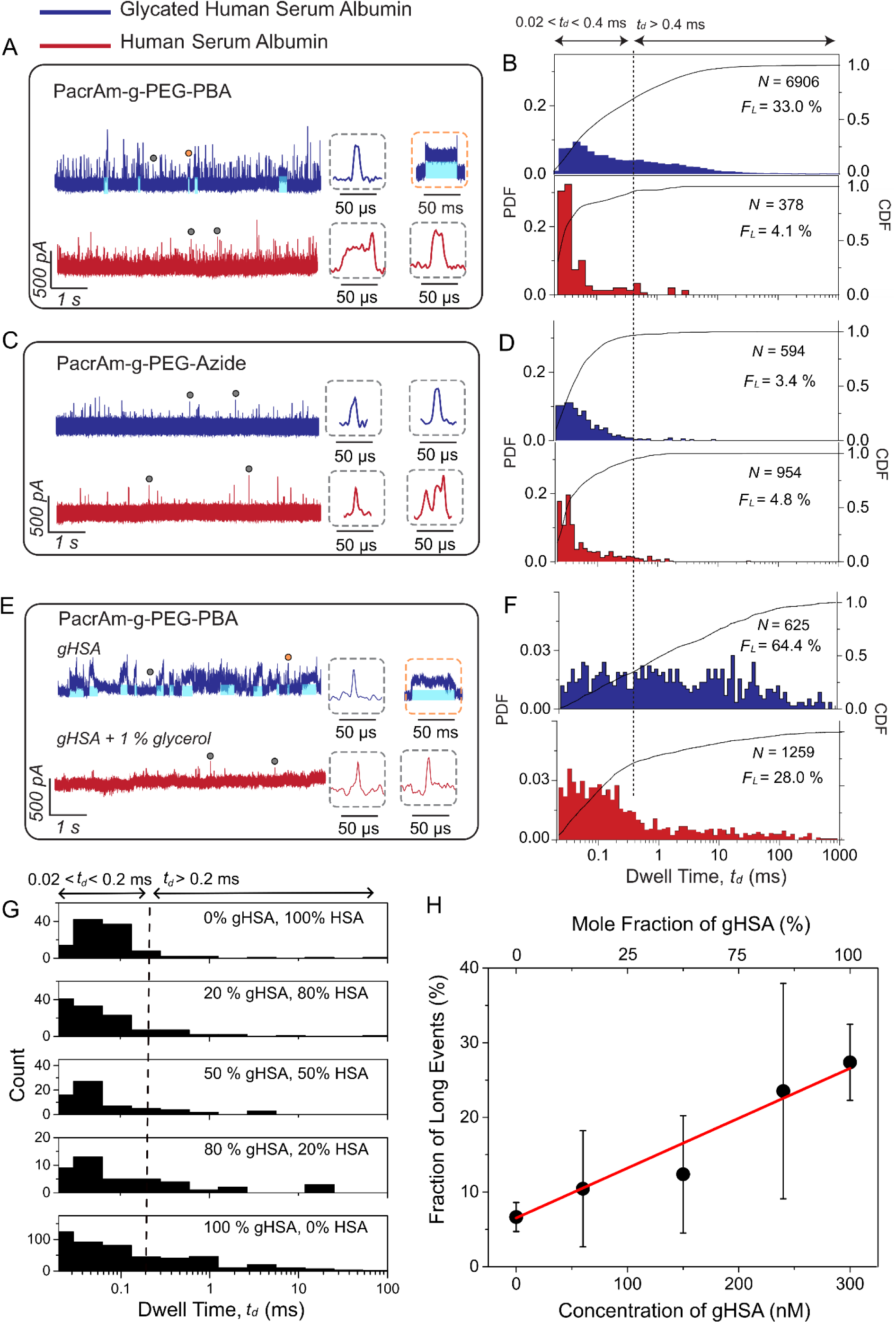
Selective trapping of glycated proteins in PBA-coated nanopores and control experiments. **A)** Representative ionic current recordings of gHSA (blue) and HSA (red) proteins obtained using a SiN_x_ nanopore with a diameter of 18 nm and coated with PAcrAm-g-PEG-PBA. Some of the trapping events (i.e., *t_d_* > 0.4 ms) in A are highlighted with blue shading. Insets display individual short resistive pulses (gray dotted boxes) and long resistive pulses (orange dotted box). **B**) Corresponding probability density functions (PDF, red and blue) and cumulative distribution functions (CDF, black curves) of dwell times for gHSA (blue) or HSA (red). *F_L_* refers to the fraction of long resistive pulses (*t_d_* > 0.4 ms) relative to all detected resistive pulses. **C**) Control experiment showing representative ionic current recordings of gHSA (blue) and HSA (red) obtained using a SiN_x_ nanopore with a diameter of 18 nm and coated with PAcrAm-g-PEG-Azide (i.e., without PBA groups). Insets display individual short resistive pulses (gray dotted boxes). **D**) Corresponding probability density functions (PDF, red and blue) and cumulative distribution functions (CDF, black curves) of dwell times for gHSA (blue) or HSA (red). **E)** Representative ionic current trace of gHSA obtained using a SiN_x_ nanopore with a diameter of 11 nm and coated with PAcrAm-g-PEG-PBA in the absence and presence of 1 % glycerol. See also in **Supplementary Figure S4. F)** Corresponding PDF and CDF of dwell times for gHSA in a nanopore with a diameter of 11 nm coated with PAcrAm-g-PEG-PBA in the absence (blue) and presence (red) of 1 % glycerol. The vertical dashed line at 400 µs marks the threshold between short and long events. Resistive pulses shorter than 20 µs were excluded due to 50 kHz low-pass filtering. **G)** Histogram of dwell times of resistive pulses from a sample containing gHSA and HSA in various ratios. The total concentration (gHSA + HSA) was fixed at 300 nM. The vertical dashed line at 200 μs marks the threshold between short and long events. The shorter dwell time threshold of 200 μs instead of 400 μs was selected because, for this experiment, a high applied potential difference of 1 V was used during the recording. **H)** Fraction of long events as a function of gHSA concentration in the mixture. A linear fit (red line) yields an *R*^2^ value of 0.99. Data were acquired using a PBA-coated nanopore with a diameter of 15 nm, sampled at 500 kHz, and filtered with a digital Gaussian low-pass filter at 50 kHz. The error bar represents the standard deviation of *F_L_* determined from resistive pulses that were detected within each 10-second recording window.

### Selectivity for binding glycated proteins in PBA-coated nanopores

To assess the specificity of trapping glycated proteins, we performed several control experiments. First, we confirmed that a PAcrAm-g-PMOXA coating without PBA groups, exhibited minimal fraction (≤ 2%) of resistive pulses with long dwell time *t_d_* > 400 µs with both gHSA and HSA (**Supplementary Figure S5**). We had previously shown that this coating is well suited to reduce non-specific interactions of proteins with nanopore wall.^21^ Next, we assessed the PAcrAm-g-PEG-Azide coating, produced during the first step of nanopore functionalization. This control coating differs from PAcrAm-g-PEG-PBA only in the absence of a PBA group with its linker. As expected, this coating exhibited predominantly (≥95%) short resistive pulses (*t_d_* ≤ 400 µs) with both gHSA and HSA proteins, indicating a low level of nonspecific protein interactions (**Figures 2C, 2D**). Nevertheless, the fraction of long dwell time events was slightly elevated (3.4%-4.8%) compared to the PMOXA coating (**Figure 2G**), suggesting that the azide groups may engage in weak non-specific interactions of proteins with the coating.^47^

In contrast to these results from nanopore coatings without PBA groups, resistive pulses with gHSA and nanopores coated with PAcrAm-g-PEG-PBA exhibited strongly increased fractions of resistive pulses with long dwell times. **Figure 2B** shows the observed wide range of dwell times spanning from 20 μs to 1 s, with the fraction of long events reaching one third of all detected events (*F_L_* = 33.0%). In contrast, for non-glycated HSA, the fraction of resistive pulses with long dwell times was similar to that of the control experiments (*F_L_* = 4.1%) as expected. The slightly higher *F_L_* value of 4% compared to the *F_L_* value of 2% for the PMOXA coating (**Supplementary Figures S5**) may reflect the presence of a small fraction of glycated HSA in the sample of HAS, which is obtained and purified from human plasma.^48^

We further investigated the specificity for trapping glycated proteins in a nanopore with a PBA coating through a competitive binding experiment, by adding glycerol—a small molecule with vicinal diols—to the protein sample. The presence of glycerol strongly reduced the fraction of resistive pulses with long dwell time (**Figures 2E, 2F, Supplementary Figure S4**), demonstrating its effective competition with gHSA for binding to the PBA coating. Note, these experiments were conducted using a smaller nanopore with a diameter of 12 nm, which increased the overall fraction of trapping events (**Figure 2F**).

To investigate the selectivity of PBA-coated nanopores for glycated proteins in a mixture containing also non-glycated proteins, we performed translocation experiments with varying concentrations of gHSA in the presence of non-glycated HSA, while keeping the total protein concentration (gHSA + HSA) constant (**Figure 2G, 2H**). As expected, the fraction of long events was proportional to the concentration or mole fraction of gHSA (*R*^2^ = 0.99). This result confirms that PBA-coated nanopores make it possible to selectively and quantitatively detect glycated proteins in a sample matrix that contains non-glycated proteins, even if the mole fraction of glycated proteins is smaller than 20%.

To explore whether the reversible trapping strategy developed here can be applied broadly to all proteins, including non-glycated proteins, by deliberately attaching moieties with vicinal diols to these proteins, we implemented a straightforward glycation step on ferritin before conducting the translocation experiments. **Figure 3** shows that non-glycated ferritin produced predominantly (99 %) short resistive pulses (*t_d_* < 0.4 ms) in a PBA-coated nanopore. Due to the strong negative charge of ferritin at the physiologic pH of 7.4, these short-lived resistive pulses could not be fully resolved from the ionic current trace. In contrast, after deliberate glycation of ferritin by incubating the protein in a buffer containing 0.5 M glucose for 12 h, the resulting glycated ferritin generated a large fraction (∼53%) of long resistive pulses.^49^ These results highlight that the protein trapping strategy introduced in **Figure 1** is broadly applicable and can be extended to natively non-glycated proteins following straightforward incubation in a glucose solution without requiring additional reagents.

**Figure 3.**
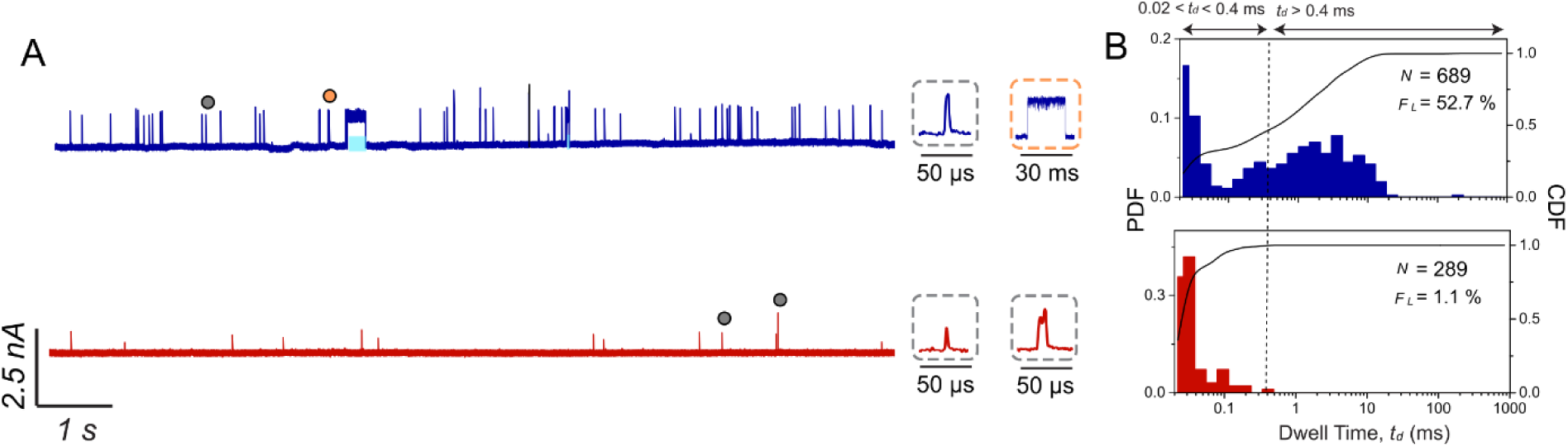
Deliberate glycation of proteins by incubation in a glucose solution makes it possible to trap previously non-glycated proteins in a PBA-coated nanopore. **A)** Representative ionic current recordings of ferritin that was deliberately glycated by incubation in a 0.5 M glucose solution with a pH of 7.5 for 12 h (blue) and non-glycated ferritin that was incubated for the same time in the same buffer without glucose (red). In both cases, we used nanopores with a diameter of 21 nm in SiN_x_ membranes coated with PAcrAm-g-PEG-PBA. Some of the trapping events (*t_d_* > 0.4 ms) in **A** are highlighted with blue shading. Insets display individual short resistive pulses (gray dotted boxes) and long resistive pulses (orange dotted box). **B**) Probability density functions (PDF, red and blue) and cumulative distribution functions (CDF, black curves) of dwell times for glycated ferritin (blue) and normal ferritin (red) in nanopores coated with PAcrAm-g-PEG-PBA. *F_L_* refers to the fraction of long resistive pulses (*t_d_* > 0.4 ms) relative to all detected events.

Together, these results demonstrate that nanopores with a polymer coating that exposes PBA groups can reliably, specifically, and transiently trap glycated proteins through reversible covalent bonds between vicinal diols and PBA groups. These transient trapping events extend the dwell times from 1-10 μs for most free translocations to 0.1 – 1 s for the longest trapping events, corresponding to five orders of magnitude.

### Controlling the most probable trapping time by the applied electric field and the choice of pH

To control the duration and frequency of individual trapping events for improved protein characterization, we examined the effects of applied voltage and pH of the recording buffer on the most probable dwell time of trapped proteins in PBA-coated nanopores. To this end, we varied the applied potential difference from −100 to −400 mV and quantified the fraction of all detected resistive pulses that were longer than 0.4 ms, *F_L_*, the dwell times of these pulses, *t_d_*, and the frequency of long events. **Figure 4A, B** shows that *F_L_* varied with applied potential, reaching a maximum of 35% at −200 mV. At higher voltages, the increased electrophoretic force accelerated protein movement, reducing the residence time of proteins inside the pore and hence the probability of reaction, and thus reducing *F_L_*. Conversely, at the lower voltage of −100 mV, the fraction of long events appears low because of the long trapping times that often exceeded 100 ms. During such long residence times, additional gHSA proteins entered the pore while previously bound proteins remained within the lumen. This effect made it difficult to identify the end of these extremely long trapping events and could lead to an unstable baseline at −100 mV.

**Figure 4.**
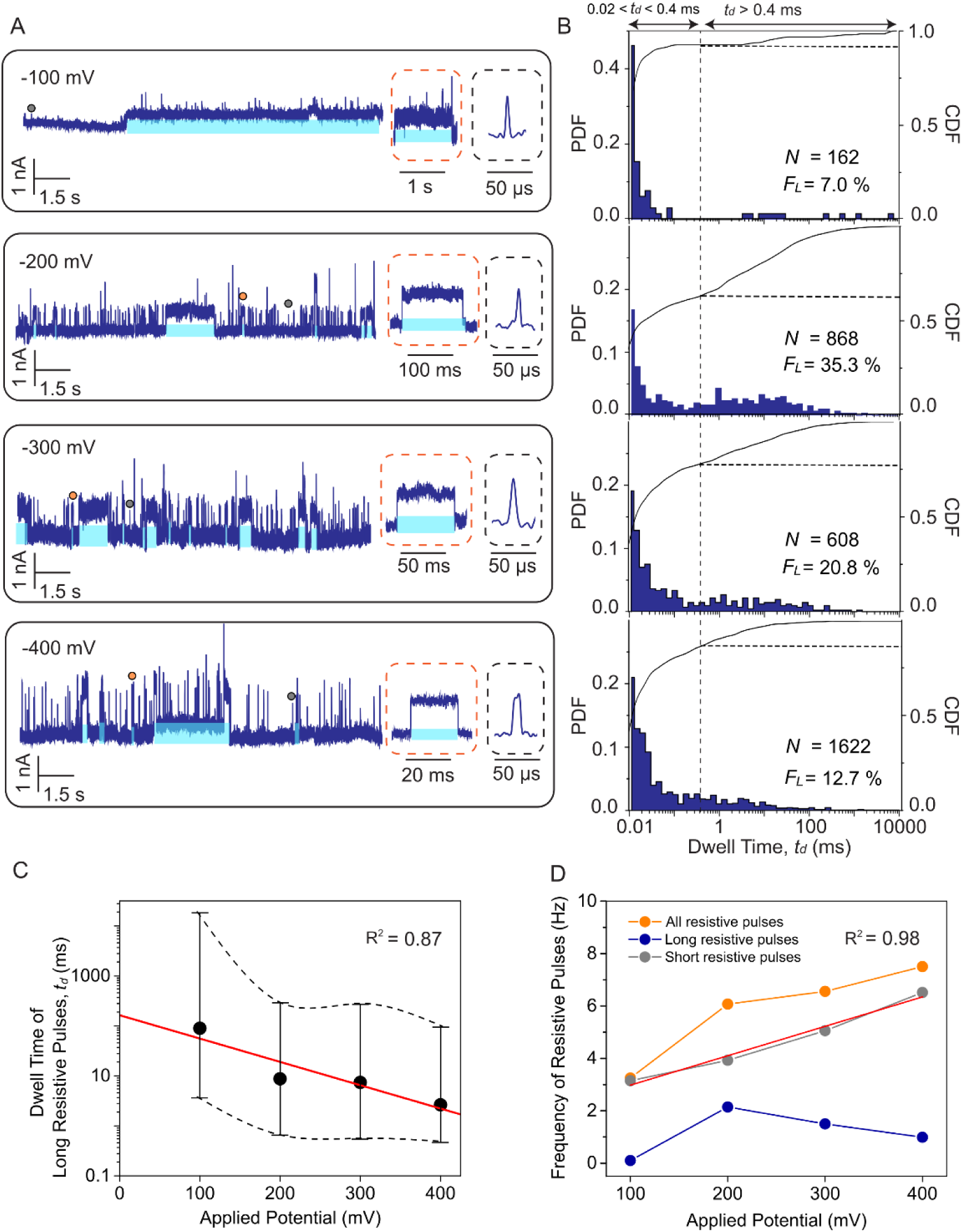
Controlling the dwell time of trapped proteins by the applied potential difference. **A)** Representative ionic current traces recorded with a PBA-coated nanopore and gHSA at different applied voltages. Insets display individual short resistive pulses (gray dashed boxes) and long resistive pulses (orange dashed boxes). Some of the trapping events (*t_d_* > 0.4 ms) in A are highlighted with blue shading. **B)** Corresponding PDF and CDF of dwell times at different applied voltages. **C)** Median dwell time of long events as a function of applied voltage. The red line represents a linear least-squares regression fit (*R*^2^ = 0.87) to the logarithm of median dwell times. Error bars indicate the 5 to 95% confidence intervals. **D)** Frequency of detected resistive pulses as a function of applied voltage for all resistive pulses (orange), long resistive pulses (*t_d_* > 0.4 ms, blue), and short resistive pulses (0.02 < *t_d_* <0.4 ms, gray). The red line represents a linear fit (*R*^2^ = 0.98) to the frequency of short events with a slope of 1.1 Hz/mV. All data were collected using the same nanopore with a diameter of 21 nm, a recording buffer containing 2 M KCl, 10 mM HEPES buffer, pH 7.5, and a sampling rate of 500 kHz and a digital Gaussian low-pass filtering with a cutoff frequency of 50 kHz.

With regard to dwell times, we observed a statistically significant (*p* = 1.05 × 10^−12^, Kruskal-Wallis test) gradual decrease in the median *t_d_* values of long events with increasing voltage (**Figure 4C**). We attribute this trend to the voltage-dependent electrophoretic force exerted on trapped gHSA with an estimated net charge of ∼15e^−^. This electric field-induced electrophoretic force presumably reduced the effective activation energy barrier required for the dissociation of the boronate ester.^50^ Based on previous work on the binding of His-tagged proteins to NTA groups in a nanopore by Wei et al.,^36^ we assumed that the median dwell time of long events decreased exponentially with voltage. Fitting **Equation S5** (**Supplementary Note S3, Figure 4C**) to the data returned a dwell time of *t_0_* = 164 ms at 0 mV applied potential corresponding to dissociation rate constant *k_off_* = 1/*t_0_* of 6.1 s^−1^ in the absence of an external electric field. This *k_off_* value is in good agreement with previously reported dissociation rates for boronate ester, which range from 2.2 to 4.3 s^−1^.^46, 51^

Finally, we analyzed the frequency of all detected resistive pulses (*t*_d_ >20 µs), long resistive pulses (*t*_d_ >400 µs), and short resistive pulses (20 µs < *t*_d_ ≤400 µs) at each applied voltage (**Figure 4D, Supplementary Note S4).** The frequency of short events increased linearly with the applied voltage, consistent with a previous analytical prediction for the capture frequency of free proteins in nanopores.^52^ In contrast, the frequency of long events exhibited a non-linear dependence on the applied potential with a maximum at −200 mV. Since the residence time of trapped proteins in the nanopore is reduced with increasing voltage. Combining the information on the frequency, fraction, and dwell time of long events, we identified −200 mV as the optimal applied potential for protein resistive pulses with transient trapping in the nanopore lumen.

Importantly, the frequency of long events remained relatively high at ∼1.0 Hz even at an applied voltage of −400 mV. In nanopore experiments, high applied voltages increase the amplitude of resistive pulses and thus have the potential to improve the signal-to-noise ratio (**Supplementary Figure S6**). In the absence of trapping, however, the benefits are undermined by reduced dwell times of resistive pulses in response to the fast electrophoretic motion of proteins at high applied potential differences (**Supplementary Figure S7**). Therefore, the reversible trapping approach used here may be beneficial for characterizing small proteins or proteins with a large net charge, as well as for other experiments that may benefit from high voltage operation.

To further control the duration and frequency of trapping individual single proteins in PBA-coated nanopores, we investigated the effect of the pH of the recording buffer on these two parameters with gHSA (**Figure 5)**. The theoretical *t_d_* values predicted from the net charge of the protein vary with pH (**Supplementary Table S1**), influencing the electrophoretic force and hence the protein’s velocity in the nanopore. The net charge also alters the residence time in the pore and hence the probability of trapping by forming a reversible covalent bond. Once the bond is formed, the electrophoretic force acting on the trapped protein affects the off-rate of the reversible bonds.^53, 54^ In addition, the reaction between PBA and vicinal diols is pH dependent; increased proton concentration at acidic pH values shifts the equilibrium toward bond dissociation (**Figure 1B**).^55, 56^ Finally, previous studies have shown that, under strong alkaline conditions (pH ≥ 9), the reaction probability between PBA groups and vicinal diols decreases due to the conversion of the reactive R-B(OH)_2_ moiety to the less reactive *R* − *B*(*OH*)^−^ moiety.^56^

**Figure 5.**
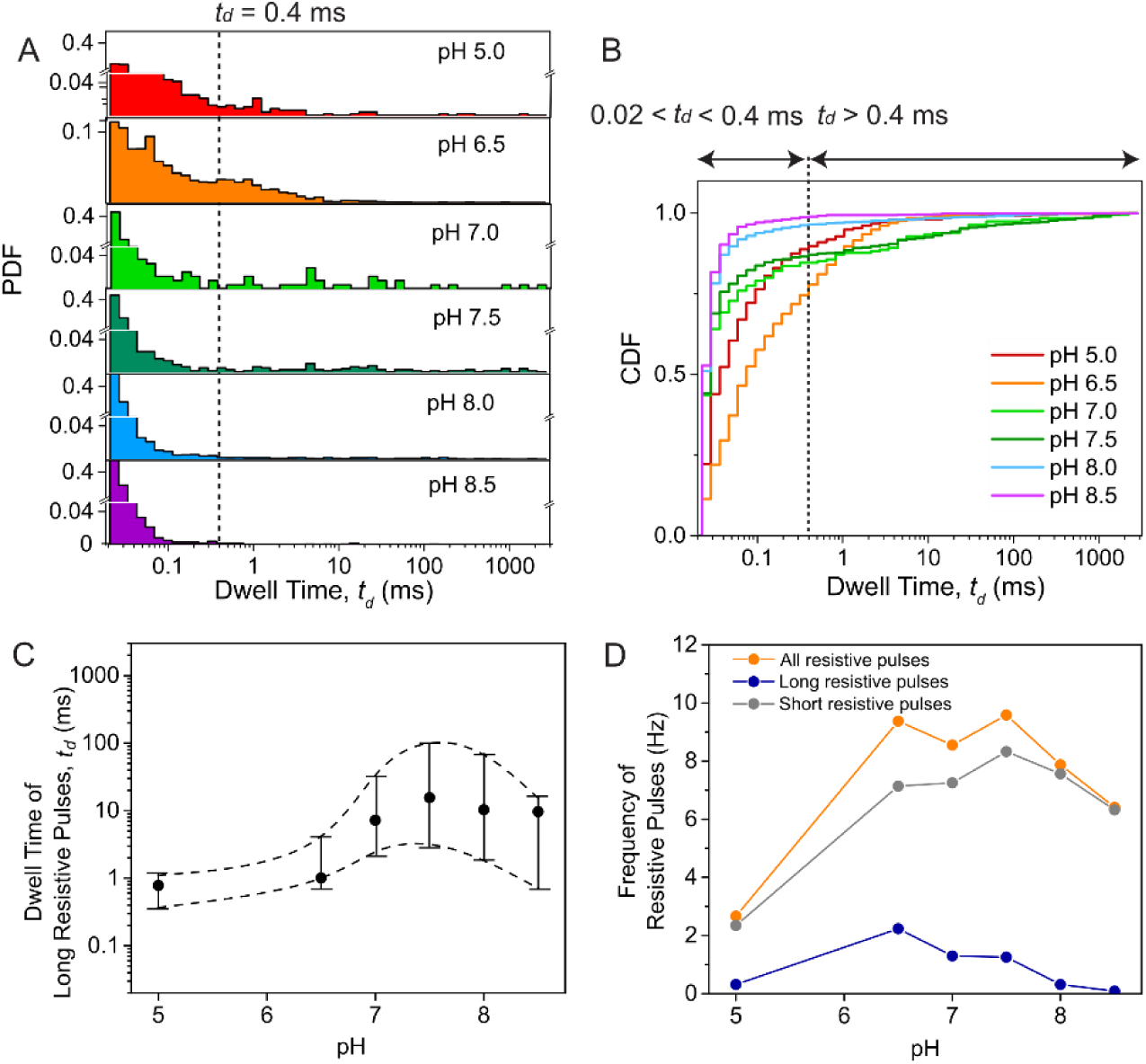
Dependence of the duration and frequency of trapping gHSA in PBA-coated nanopores on the pH value of the recording buffer. **A**) Distribution of dwell times at different pH values of the recording buffer. Recording buffers contained 2 M KCl with 10 mM MES for pH 5.0, 10 mM Bis-Tris for pH 6.0 - 6.5, 10 mM HEPES for pH 7.0 - 8.0, or 10 mM AMPSO for pH 8.5. **B)** Cumulative density functions (CDF) of dwell times at various pH values. **C)** Median dwell time of long resistive pulses as a function of the pH of the recording buffer. Whiskers represent 5% and 95% confidence intervals. **D)** Frequency of resistive pulses as a function of the pH of the recording buffer of all detected resistive pulses (orange), long resistive pulses (blue), and short resistive pulses (gray). Data were collected using the same nanopore with a diameter of 19 nm for all experiments at a sampling rate of 500kHz under an applied voltage of −200 mV and a digital Gaussian low-pass filter with a cutoff frequency of 50 kHz.

**Figure 5** shows the fraction of long resistive pulses, their dwell times, and the frequency of protein trapping across a range of pH values of recording buffer ranging from 5.0 to 8.5. The fraction of long resistive pulses peaked at ∼22.1 % at pH 6.5, as shown in the CDF (**Figure 5B)**, and remained relatively high at neutral pH (∼15% at pH 7.0 and ∼13% at pH 7.5). The median dwell time of long pulses increased tenfold with increasing pH (*p* = 1.3 × 10^−14^ Kruskal-Wallis test), reaching ∼10 ms at neutral and slightly basic pH values (7.0 - 8.5, **Figure 5C**). On the other hand, the frequency of detectable long resistive pulses exhibited a maximum of 2.3 Hz at pH 6.5. At slightly basic pH (8.0 - 8.5), the frequency of long resistive pulses was smaller than 2% with approximately 98% of short resistive pulses (**Figures 5B, 5D**). This observation suggests that either the increased net charge of the protein at basic pH levels reduced the residence time of gHSA in the pore and hence reduced the probability of forming a reversible covalent bond, or that these alkaline pH values already reduced the propensity for reaction.^56^ It is also possible that both mechanisms contributed to the reduced frequency of long resistive pulses. Considering these results, we identified neutral pH (7.0 - 7.5) as optimal for experiments to achieve relatively frequent and transient protein trapping in the nanopore lumen. If, however, the most important parameter is to extend trapping time for as long as possible, then pH 7.5 and pH 8.0 resulted in median trapping times of up to 100 ms when −200 mV was applied and up to ∼1 s at −100 mV applied potential (**Figure 5C**).

### Determination of the volume and shape of single glycated proteins using PBA-coated nanopores

To evaluate the performance of PBA-coated nanopores for determining the volume and shape of single trapped proteins, we analyzed gHSA (MW = 66.5 kDa, pI = 4.4) and three other naturally glycated proteins: human hemoglobin A1c (HbA1c, MW = 64.5 kDa, pI = 6.9), human immunoglobulin G (IgG, MW = 150.0 kDa, pI = 7.3), and human thyroglobulin (Tg, MW = 660.0 kDa, pI = 4.5) (**Figure 6A**).

**Figure 6.**
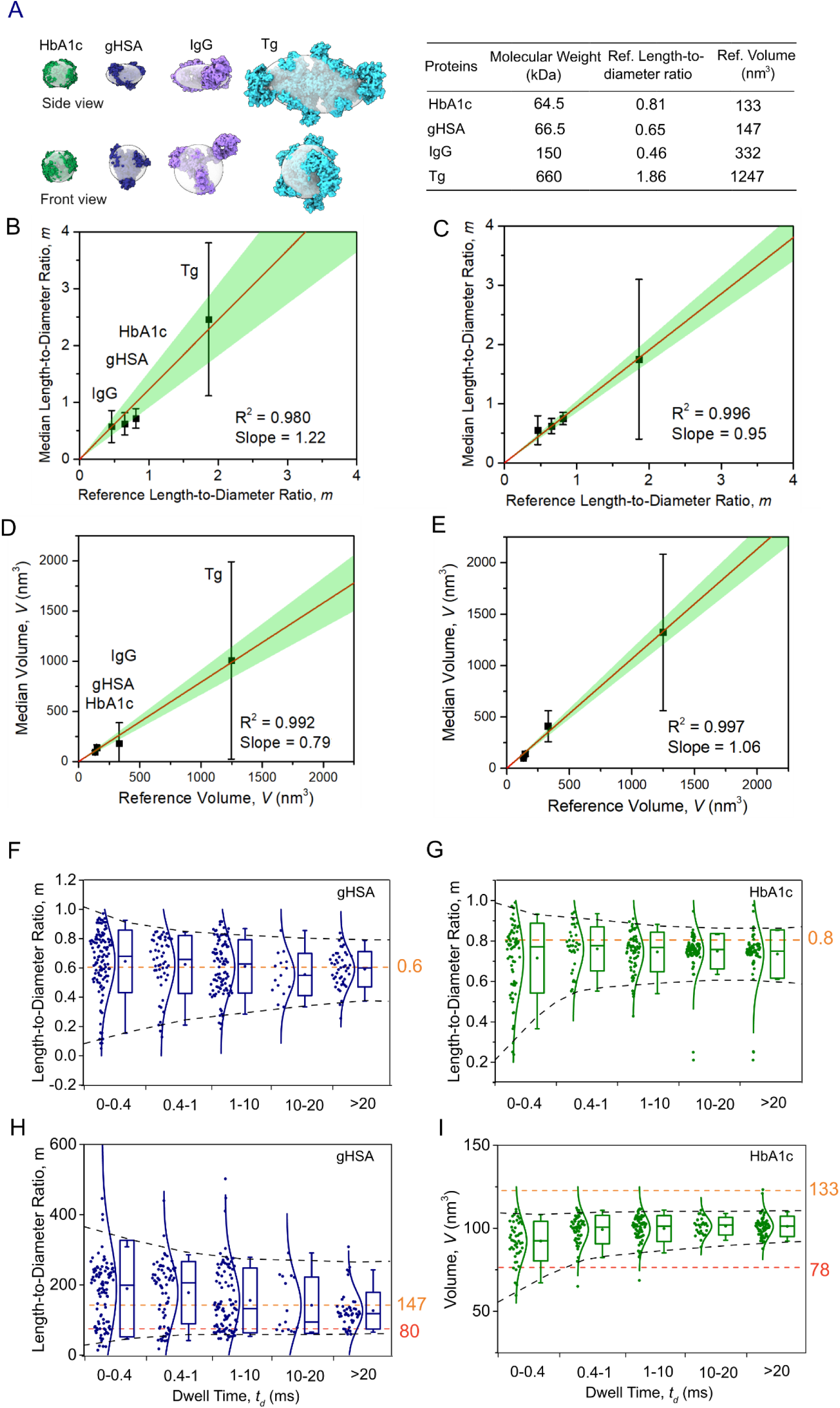
Accuracy of nanopore-based volume and shape determination of single proteins and effect of trapping time. **A**: Atomic structures of the four test proteins alongside their theoretically calculated reference ellipsoids (in transparent gray). Green – HbA1c, (PDB: 1A3N); blue – gHSA, (PDB: 1AO6); purple – IgG, (PDB: 1HZH); cyan – Tg, (PDB: 6SCJ). **B, C, D, E:** Median values of excluded volume *V*, and length-to-diameter ratio *m* plotted as a function of the expected reference values (see Methods) for individual proteins. Panels **B** and **D** correspond to the analysis of short events (150 µs < *t_d_* < 400 µs), panels C and E to the analysis of long events (*t_d_* > 400 µs). Whiskers represent the 5% and 95% confidence intervals of experimental estimates. The red line shows a linear least squares regression fit constrained to a zero intercept, with the green shaded area indicating the 95% confidence interval for the fit. **F, G**: Estimated length-to-diameter ratio *m* as a function of trapping time for (**F**) gHSA and (**G**) HbA1c, demonstrating the improvement of shape estimates with increasing trapping time. Orange dashed lines indicate reference length-to-diameter values (see Methods). Black dashed lines indicate the 5% and 95% confidence intervals. **H, I**. Estimated volume *V* as a function of trapping time for **(H)** gHSA or **(I)** HbA1c. Orange dashed lines indicate reference volume estimated using solvent accessible surface (see Methods). Red dashed lines indicate the volume estimated using molecular weight.^57^ For both proteins, estimates of the shape, and to a smaller extent, estimates of the volume improve with increasing residence time.

Since accurate estimation of protein volume, *V,* depends on the geometry of the nanopore lumen, we first calibrated all nanopores using the spherical protein streptavidin (length-to-diameter ratio *m* ≈ 1) to confirm that the analysis results in a round shape and to determine the effective length of the nanopore (**Supplementary Note S5**).^10, 21^ Proceeding only with nanopores that resulted in an *m* value close to 1.0 for streptavidin, we determined first the effective length of the nanopore based on the resistive pulse from streptavidin translocation and then the volume, *V*, and shape, *m*, of these four proteins using a previously described approach.^10^ **Figure 6** shows that by analyzing all resistive pulses with *t_d_* values exceeding 150 µs, we obtained median volume and shape values that closely matched reference values derived from atomic structures. These results demonstrate the accuracy of PBA-coated nanopores for measuring protein volume and shape (see also **Supplementary Table S2, Supplementary Figures S8A, B**). The distributions of dwell times for each protein revealed populations of long events (*t_d_* > 400 µs), indicating trapping by reversible covalent bonds of these four glycated proteins and PBA groups in the nanopore (**Supplementary Figure S8C)**. These results suggest broad applicability of the approach to glycated or glycosylation proteins. To assess the benefits of long-lasting trapping events for quantitative volume and shape analysis, we separated short (150 µs < *t_d_* ≤ 400 µs) and long events (*t_d_* ≥ 400 µs) and calculated the excluded volume, *V,* and length-to-diameter ratio, *m,* separately for each group (**Figures 6B-E)**. Median estimates based on long events deviated on average from the reference values by only +15.1% with respect to volume and −9.4% with respect to *m* value, representing an improvement in accuracy by ∼10% compared to short events, which deviated from reference values by −25.1 % with respect to volume and +17.8 % with respect to shape (**Supplementary Table S3**). We attribute this improved accuracy to the improved sampling and statistics that result from long residence times of the trapped proteins in the nanopore. Consistent with this improvement, the correlation between experimental and reference values for both shape and volume improved for long events, as reflected by higher Pearson’s correlation coefficients (**Figures 6B-E**).

One important aspect that the approach of reversible trapping introduced here makes possible to explore is the minimum required trapping time to obtain an accurate estimation of the volume and shape of single trapped proteins in the nanopore. **Figures 6F, G, H, and I** show that the optimal residence time in the pore is between 1 and 20 ms. Longer tapping times offered no significant further improvement of characterization, while shorter times do increase the uncertainty of shape and volume determination, presumably due to incomplete sampling of all representative protein orientations in the pore, and insufficient statistics for these short events.

These results demonstrate that the approach of transient covalent trapping of a protein in a nanopore with a flexible tether introduced here does not adversely affect the analysis of volume and shape. On the contrary, estimates of these two parameters from individual resistive pulses become more reliable as dwell times reach between 1 and 20 ms. Trapping times longer than 20 ms are not desirable as they reduce the throughput of analysis and increase the risk of trapping more than 1 protein in the pore, making shape determination impossible.

## Conclusion

This work introduces a versatile and straightforward-to-use polymer coating for nanopores in SiNx membranes based on commercially available PAcrAm-g-PEG-Azide molecules. This coating serves as an inert layer that strongly reduces non-specific protein adsorption to the nanopore walls (**Figures 2C,D**). A one-step copper-free click reaction makes it possible to convert the pending surface azide groups into covalently attached PBA groups. These PBA groups enable selective transient trapping of glycated proteins by exploiting a reversible covalent bond between PBA and vicinal diols on glycated proteins. Moreover, this PBA-activated coating remains resistant to non-glycated proteins (**Figure 2**). With minor modifications, this approach could also be used to incorporate other active groups, such as nitrilotriacetic acid (NTA) for His-tag affinity, biotin for streptavidin recognition, or maleimide for cysteine-specific coupling onto the nanopore surface.

This study establishes the broad applicability of reversible covalent trapping of proteins to slow down their translocation through nanopores. Straightforward incubation of non-glycated proteins in a glucose solution leads to their glycation and extends the approach to all proteins with accessible amine groups. This versatility highlights the potential of reversible covalent trapping as a universal strategy for enhancing temporal resolution in nanopore sensing and paves the way for more accurate analysis of complex protein systems. Importantly, the covalent bond for trapping occurs between two small chemical moieties, PBA and a carbohydrate with vicinal diols such as glucose. These small binding partners do not significantly change the size and shape of the trapped proteins while providing most probable trapping times of tens of milliseconds. This aspect is critical for single protein characterization.

We demonstrate that the choice of applied voltage and the pH value of the recording buffer make it possible to control and fine-tune the probability and duration of protein trapping. We show that resistive pulses with a duration between 1 and 20 ms provide close to maximum accuracy of protein volume and shape estimation with an ellipsoidal model (**Figure 6, Supplementary Table S3**). Finally, the ability to prolong the interrogation of individual proteins may open the door for analyzing molecular shape beyond the simple ellipsoid approximation,^58^ paving the way for more advanced nanopore-based studies of protein conformation.

## Materials and Methods

### Materials

Poly(acrylamide)-g-PEG-Azide^21, 32^ (M.W. ∼68 kDa) and Poly(acrylamide)-g-PEG-PMOXA (M.W. ∼68 kDa), designed to minimize protein adhesion, were purchased from SuSoS AG, Switzerland. DBCO-PEG4-NHS (cat. n. BP-22288) was purchased from BroadPharm, USA. 4-(Aminomethyl) phenylboronic acid hydrochloride (cat. n. H52855-03) was purchased from Alfa Aesar, USA. Glycated human serum albumin (cat. n. A8301-25MG), human serum albumin (cat. n. A1653-1G), and human thyroglobulin (cat. n. T6830-1MG) were purchased from Sigma, USA. Human HbA1c (cat. n. PRO-299) was purchased from ProSpec, Israel. Immunoglobulin G (cat. n. 340-21) was purchased from Lee BioSolutions, USA. Ferritin from human spleen (cat. n. 9007-73-2) was purchased from Sigma-Aldrich, USA.

Ultra-0.5 centrifugal filter units (cat. n. UFC510024) were purchased from Sigma, USA. Syringe filters with 13 mm diameter and 220 nm pore size (cat. n. SF1303-1) were purchased from BGB Analytik, Switzerland. Anotop syringe filters with 10 mm diameter and 20 nm pore size (cat. n. 6809-1002) were purchased from Fisher Scientific. SiN_x_ chips with a single nanopore with diameters of 15 nm, 20 nm, or 25 nm in a free-standing SiN_x_ membrane were made by helium-focused ion beam by Norcada Inc., Canada.

### Preparation of PBA with DBCO activation

We dissolved 25 mg DBCO-PEG_4_-NHS or DBCO-PEG_12_-NHS in 1 mL of DMSO, yielding a concentration of 25 mg/mL. One molar equivalent of Amino-Phenylboronic acid and 1.5 molar equivalents of Triethylamine were mixed and then added to the DMSO solution. The reaction mixture was stirred at 60 rpm for 12 h at room temperature under a nitrogen atmosphere. The reaction product (DBCO-PEG-PBA) was analyzed for purity by NMR and HPLC, aliquoted into 50 µL portions, and stored at −80 °C. These analyses revealed a purity of > 90%.

### Coating of Nanopore Chips

Coating solution A contained 0.1 mg/mL PAcrAm-g-PEG-Azide in 1 mM HEPES buffer, pH 7.4. Coating solution B contained 1.2 mg/mL DBCO-PEG4-Phenylboronic acid (PhB(OH)_2_) in 10 mM HEPES pH 7.4, 160 mM NaCl. Both solutions were filtered using a 220 nm syringe filter.

SiN_x_ chips were cleaned by oxygen plasma using a plasma cleaner (Diener electronic, Germany) operated at 30 % power and 0.3 mbar O_2_ for 30 s on each side. The chips were immersed in coating solution A and incubated for 60 min at room temperature. After incubation, the chips were rinsed three times with ultrapure water and dried under a stream of nitrogen. The dried chips were subsequently immersed in coating solution B for 60 min at room temperature, followed by rinsing with ultrapure water and drying with a stream of nitrogen.

To verify selective binding of glycated proteins to PBA groups on the inner wall of the nanopore, we compared resistive pulses in the presence of gHSA or HSA through nanopores with different surface coatings: One with PBA group, namely, PAcrAm-g-PEG-PBA (Figures 2A, 2B), PAcrAm-g-PEG-Azide (with azide functional groups, Figures 2C, 2D), and PAcrAm-g-PMOXA (Supplementary Figure S6). These experiments were conducted using nanopores with a diameter of 18 nm, a recording buffer with pH 7.5, and an applied voltage of −200 mV.

### Deliberate glycation of proteins

Ferritin from human spleen (Sigma-Aldrich) was dissolved in phosphate-buffered saline (PBS). D-(+)-glucose (Sigma-Aldrich) was dissolved at a 0.5 M concentration in PBS and filtered through a 0.22 μm membrane. All reagents were of analytical grade. The ferritin concentration was adjusted to 1 mg/mL in PBS, and we then added glucose spontaneously to a final concentration of 0.5 M in a total reaction volume of 0.1 mL. Glycation occurred by incubation at 37 °C for 24 h (150 rpm).^49^ After incubation, unreacted glucose was removed by dialysis using a 10 kDa MWCO membrane against 150 mL PBS at 4 °C. The dialysis buffer was replaced every 2 h for a total of two changes.

### Recordings of Electrical Resistance

Ionic currents through the nanopores were recorded using a patch-clamp amplifier (AxonPatch 200B, Molecular Devices, UK) with Ag/AgCl pellet electrodes (Warner Instruments, USA) under voltage clamp mode with a 100 kHz Bessel lowpass filter. The data was acquired using a data acquisition card, NI PCI 6281 (National Instruments, USA), with a sampling rate of 500 kHz. Custom-built control software based on the Python version of the NI-DAQmx library (National Instruments, USA) was used to collect data. To optimize the signal-to-noise ratio, we selected nanopore diameters based on the molecular weight of proteins as described earlier.^12^ To this end, we used nanopores with the following diameters: 19 nm for HbA1c, 22 nm for gHSA, 25 nm for IgG, and 35 nm for Tg. All experiments were conducted in a recording buffer with a pH of 7.5 at an applied voltage of −200 mV.

### Preparation of Protein Solutions

Nanopore experiments were conducted in a recording buffer containing 2 M KCl, 10 mM HEPES, pH 7.5, which was filtered using a 20 nm syringe filter prior to experiments. All tested proteins, except HbA1c, were used at a final concentration of 100 nM in the nanopore experiments. HbA1c was used at a final concentration of 20 nM.

Three proteins—gHSA, HSA, and HbA1c—required purification to obtain monomeric solutions before use in the nanopore experiments. For protein purification, we used either size exclusion HPLC or centrifugation in a filter unit (MWCO 100 kDa, Amicon), with both methods yielding similar purity. In the HPLC method, proteins were diluted to a concentration of 1 mg/mL in PBS buffer and separated using an Agilent SEC3 HPLC column (300 mm length, 300 nm pore size, 4.6 mm internal diameter, Agilent Technologies, USA). The running buffer consisted of PBS supplemented with 1 mM EDTA, and the flow rate was 0.3 mL/min. The monomer fraction of proteins was collected, aliquoted, and stored at −80 °C. Before use, an aliquot was thawed, and the buffer was exchanged to recording buffer using a centrifugal filter unit with a 50 kDa MWCO (Amicon).

For purification via centrifugation, proteins were diluted to 1 mg/mL in the recording buffer and centrifuged in a filter unit with a 100 kDa NMWCO at 12298 g for 5 min. The collected permeate solution contained monomeric protein at a typical concentration of 0.5–1 µM, which was used as stock for nanopore experiments.

### Data Analysis

We used home-built software written using C++, which is called Visual Nanopore, to identify and analyze individual resistive pulses. Visual Nanopore uses a two-sliding window algorithm^59^ to identify the resistive pulses. Protein shape and volume were analyzed only for resistive pulses lasting longer than 150 µs using a home-built MATLAB program, as previously described^60^ and briefly explained in **Supplementary Note S5**. We excluded resistive pulses from the analysis if their determined *ΔI_min_* values were smaller than 5 times the standard deviation of the open pore current *I_0_*.

### Handling of Atomic Structures

We used ChimeraX^61^ to visualize the atomic structures of proteins, and we estimated the net charge of proteins using the APBS calculation.^62^ To estimate the reference ellipsoid parameters, we used Minimum Volume Enclosing Ellipsoids (MVEE)^63^ in order to fit the parameters of each axis that determines the shape of the ellipsoids. Protein volumes were then calculated based on the solvent accessible surfaces^64^, employing a custom depth-first searching algorithm with a water probe of diameter 0.28 nm. The algorithm was written with C++, and we provide an executable file to perform the shape and volume fitting from a PDB file.

## Acknowledgments

M.M. acknowledges financial support from the Swiss National Science Foundation (Grant number: 200020_197239) and from the Adolphe Merkle Foundation. Y.J.L. and S.A. were supported by grant 200020_197239 and acknowledge the Adolphe Merkle Foundation for support. Zhenghao Dong and Youwei Ma for their help on polymer synthesis, as well as Anasua Mukhopadhyay and Andela Vracar for their help on protein purification.

