## Supplementary Information for "Selective and Controllable Trapping of Single Proteins in Nanopores using Reversible Covalent Bonds"


### Supplementary Note S1: Characterization of polymer coatings for nanopores

In order to estimate the volume and shape of single proteins from resistive pulse recordings, the electric field inside the nanopore must be as uniform as possible.[1] To this end, the shape of the nanopore should approach as well as possible an ideal cylinder. The dimensions of a perfectly cylindrical nanopore can be determined from the ionic current using Equation S1:

$$G = \frac{\sigma}{\frac{l_p}{\pi {r_p}^2} + \frac{1}{2r_p}} \quad (S1)$$

Here, $G$ ($\Omega^{-1}$) is the conductrance of the nanopore measured from the *I-V* curve, $\sigma$ ($\Omega^{-1}m^{-1}$) is the conductivity of the electrolyte solution, $l_p$ (m) is the length of the nanopore channel, and $r_p$ (m) is the radius of the nanopore.

After coating the nanopore with a polymer, both the length and radius change by the effective thickness of the coating layer, $l_c$. We determined the thickness of the coating experimentally from the measured conductance, using Equation S2:

$$G = \frac{\sigma}{\frac{l_p + 2 * l_c}{\pi (r_p - l_c)^2} + \frac{1}{2(r_p - l_c)}} \quad (S2)$$

### Supplementary Note S2: Glucose adsorption fitted with a dual-site Langmuir model

Glucose contains two vicinal diol motifs (C1–C2 and C5–C6)[2] that can form reversible boronate ester bonds with PBA groups on the PAcrAm-g-PEG-PBA coating. As a result, the adsorption process is better described by a two-site Langmuir isotherm:

$$y = \frac{B_1 x}{k_1 + x} + \frac{B_2 x}{k_2 + x} \quad (S3)$$

Here, $B_1$ and $B_2$ represent the maximum adsorption capacities of the two binding modes, while $k_1$ and $k_2$ represent their respective dissociation constants, reflecting binding affinities.

### Supplementary Note S3: Calculation of dissociation rate constant for the reaction between PBA functional group and gHSA

When a protein is trapped by a reversible covalent bond, the bond will be affected by the electrophoretic force acting on the trapped protein in the strong electric field inside the nanopore.[3] Therefore, the change in Gibbs Free Energy, $\Delta G$, for the bond breaking is directly influenced by the external force.[4] The rate constant for the dissociation reaction, *k,* is hence related to the Arrhenius equation in the following way:

$$k = Ae^{\Delta G - Eq\alpha} \quad (S4)$$

Here, $A$ is the pre-exponential (Arrhenius) factor, $\Delta G$ is the Gibbs energy of activation in the absence of an external electric field, $E$ is an external electric field, $q$ is the net charge of the protein with its linkers, and $\alpha$ is a coefficient related to the length of nanopore channels.[5]

Taking the natural logarithm of **Equation S4** yields a linear relationship (**Equation S5**), which makes it possible to estimate the dissociation rate constant $k_0$ at 0 mV applied potential through least-squares fitting of the experimental data shown in **Figure 4C**:

$$lnk = lnk_0 - Eq\alpha \quad (S5)$$

### Supplementary Note S4: Calculation of the frequency of events

Since the frequency of resistive pulses may vary over time, we used instantaneous frequency to compute the frequency of long and short events of individual resistive pulses:

$$f_i = \frac{2}{t_{i+1} - t_{i-1}} \quad (S6)$$

Where $f_i$ is the frequency of $i^{th}$ event, and $t_{i-1}$ and $t_{i+1}$ are the start times of the $(i-1)^{th}$ and $(i+1)^{th}$ events. The total event frequency was determined as a mean value across all instantaneous frequencies.

### Supplementary Note S5: Determination of the effective nanopore length using resistive pulses from a spherical protein

We determined the effective nanopore length, $l_{p,eff}$, with the spherical protein streptavidin whose molecular shape can be approximated as a sphere with a length-to-diameter ratio, $m$ = 1, electrical shape factor, $\gamma$ = 1.5, and volume, $V_{SA}$ = 101 nm$^3$.[1, 6]

To characterize each nanopore, we recorded approximately one hundred resistive pulses with streptavidin ($^{\Delta I}/_{I_0}$). For quantitative analysis, we used the nanopore diameter $d_p$ as provided by TEM image and derived the effective length of the nanopores, $l_{p,eff}$, from Equation 1 as follows:

$$l_{p,eff} = \frac{6V_{SA}I_0}{\pi d_p^2 \Delta I} - 0.8d_p \quad (S7)$$

### Supplementary Note S6: Estimation of protein shape and volume

We detected resistive pulses by the so-called two sliding windows algorithm, which has been developed from the threshold detection algorithm.[7] We use the analysis of individual resistive pulses to calculate the shape and volume of the proteins based on previous work from our group by Houghtaling et al.[8] The rotation of non-spherical proteins while they move through the nanopore changes their orientation relative to the electric field and modulates the recorded ionic current blockade, $\Delta I$ as a function of the electrical shape factor, $\gamma$. The probability density function of $\Delta I$ can be described as **Equation S8** for oblate and **Equation S9** for prolate ellipsoids:

$$P(\Delta I_{\gamma}) = \frac{1}{A}\cosh\left(\frac{E\mu\left(\sqrt{\frac{\Delta I - \Delta I_{min}}{\Delta I_{max} - \Delta I_{min}}}\right)}{k_B T}\right)\frac{1}{\pi\sqrt{(\Delta I - \Delta I_{min})(\Delta I_{max} - \Delta I)}} \quad (S8)$$

$$P(\Delta I_{\gamma}) = \frac{1}{A}\cosh\left(\frac{E\mu\left(\sqrt{\frac{\Delta I - \Delta I_{max}}{\Delta I_{min} - \Delta I_{max}}}\right)}{k_B T}\right)\frac{1}{\pi\sqrt{(\Delta I - \Delta I_{min})(\Delta I_{max} - \Delta I)}} \quad (S9)$$

Here, $P(\Delta I_{\gamma})$ is the probability density distribution of $\Delta I$, which is dependent on the orientation-dependent electrical shape factor, $\gamma$. $E$ is the electric field, $\mu$ is the dipole moment, $A$ is a normalization constant for integration, and $\Delta I_{min}$ and $\Delta I_{max}$ are the minimum and maximum of ionic current from the respective resistive pulses as a response to the rotation of a single protein during its translocation.

This probability distribution does not account for noise in the current recording, so we convolve this U-shaped distribution with a standard Gaussian noise distribution function as shown in **Equation S10**:

$$P(\Delta I_{\sigma}) = \frac{1}{\sqrt{2\pi\sigma^2}} e^{-\frac{\Delta I_{\sigma}{}^2}{2\sigma^2}} \quad (S10)$$

To this end we convolve the U-shaped probability distribution with standard Gaussian noise distribution to describe the probability of the blockade of current, $\Delta I$.

$$P(\Delta I) = P(\Delta I_s) * P(\Delta I_\sigma) \quad (S11)$$

To carry out this convolution, we employed the *lsqcurvefit* function by MATLAB with four parameters that we initialized to follows: the protein permanent dipole moment, $\mu$ = 550 D, $\Delta I_{min}$ and $\Delta I_{max}$ are 5$^{th}$ and 95$^{th}$ percentiles of the current blockade, the $\sigma$ is greater than, or equal to, the standard deviation of baseline noise. The length-to-diameter ratio, *m*, of the proteins was calculated by the following equations.

*For an oblate-shaped protein:*

$$\frac{\Delta I_{max}}{\Delta I_{min}} = \left(\frac{m \cdot \cos^{-1}(m)}{(1-m^2)^{1.5}} - \frac{m^2}{1-m^2}\right)^{-1} - 0.5 \quad (S12)$$

*For a prolate-shaped protein:*

$$\frac{\Delta I_{min}}{\Delta I_{max}} = \left(\frac{m^2}{m^2-1} - \frac{m \cdot \cos^{-1}(m)}{(m^2-1)^{1.5}}\right)^{-1} - 0.5 \quad (S13)$$

Here the $\Delta I_{min}$, and $\Delta I_{max}$ are determined by fitting **Equations S12** or **S13** to the probability density distribution function of $\Delta I$. $\gamma_{\parallel}$ describes the values of electrical shape factor of the ellipsoid when its singleton axis is aligned parallel to the electric field, ***E***. $\gamma_{\perp}$ describes the singleton axis aligned perpendicular to the electric field, ***E***. **Equations S14** or **S15** describe the calculation of $\gamma_{\parallel}$ and $\gamma_{\perp}$ for oblate and prolate, respectively.

$$\gamma_{\parallel} = \frac{\Delta I_{max}}{\Delta I_{min}} + 0.5 \quad (S14)$$

$$\gamma_{\perp} = \frac{\Delta I_{min}}{\Delta I_{max}} + 0.5 \quad (S15)$$

Perfect spheres always have a constant electrical shape factor of 1.5. We determine the volume of protein, *V*, by the theory of Maxwell's derivation for translocating particle trace[6]:

$$\frac{\Delta I}{I_0} = -\frac{4V\gamma}{\pi d_p^2\left(l_p + 0.8 d_p\right)}\left(\frac{1}{1 - 0.8\left(\frac{d_m}{d_p}\right)^3}\right) \quad (S16)$$

Here, $V$ is the volume of the protein, $\gamma$ is the electrical shape factor, $d_p$ is the diameter of the nanopore, $l_p$ is the length of the nanopore channel, and $d_m$ is the diameter of the protein. To enable rotation of proteins during their translocation, we used nanopores with diameters that were at least twice the longest dimension of all tested proteins.

**Supplementary Table S1. Theoretically estimated net charge and corresponding theoretically predicted dwell time of translocation of gHSA as a function of pH.**

| pH | Theoretical charge ($e^-$)[a] | Theoretical dwell time (μs)[b] |
|---|---|---|
| 5.0 | +4.0 | 5.6 |
| 6.0 | -2.9 | 5.1 |
| 6.5 | -6.0 | 4.0 |
| 7.0 | -8.0 | 3.2 |
| 7.5 | -15.0 | 2.1 |
| 8.0 | -16.0 | 2 |
| 8.5 | -18.0 | 1.8 |

*(a) Net charge of human serum albumin is calculated from the crystal structure using APBS&PDB2PQR web service (https://server.poissonboltzmann.org/). This calculation did not account for glycation.*

*(b) Theoretical dwell time represents the most probable values from the first passage time distribution of free translocation.*[9] Using the following equation:

$$PDF = \frac{l}{\sqrt{4\pi D t^3}} e^{\frac{-\left(l-\frac{qDE}{k_B T}t\right)^2}{4Dt}}$$

with the following parameters: length of nanopore, $l$: 30 nm, electrical field, $E$: 3.3e$^{6}$ V/m, diffusion coefficient of proteins, $D$: 1e$^{-11}$ m$^2$/s.

**Supplementary Table S2. Comparison of molecular weight, isoelectric point, length-to-diameter ratio, and excluded volume of four test proteins.**

| Protein | Molecular Weight (kDa) [a] | pI [b] | Length-to-diameter ratio [c] | Volume ($nm^3$) [d] | Volume ($nm^3$) [e] |
|---|---|---|---|---|---|
| gHSA | 66.5 | 4.7 | 0.65 | 147 | 80 |
| HbA1c | 64.5 | 6.9 | 0.81 | 133 | 78 |
| IgG | 150 | 7.3 | 0.46 | 332 | 190 |
| Tg | 660 | 4.5 | 1.86 | 1247 | 790 |

*(a) Obtained from the protein data bank (https://www.rcsb.org).*

*(b) Estimated from the amino sequence using web service (https://www.protpi.ch/Calculator/ProteinTool).*

*(c) Calculated from the crystal structure using Minimum Volume Enclosing Ellipsoids (MVEE) fitting.*[10]

*(d) Calculated from the crystal structure using Solvent Accessible Volume (SAV) with a diameter of probe of 0.28 nm.*[11]

*(e) Calculated from the molecular weight*[12] *with following equation:*

$$V = \frac{4}{3}\pi(0.066\sqrt[3]{MV})^3$$

**Supplementary Table S3. Accuracy of volume and shape estimation of individual proteins using resistive pulse data from short and long resistive pulses.**

| | gHSA | HbA1c | IgG | Tg | Average absolute value |
|---|---|---|---|---|---|
| Ref. *V* | 147 | 133 | 332 | 1248 | |
| 150 µs< $t_d$ <400 µs | 139 ± 27 | 93 ± 14 | 180 ± 205 | 1006 ± 983 | |
| Percent deviation (%) | -5.3 | -30.1 | -45.6 | -19.4 | 25.1 ± 42.3 |
| $t_d$ > 400 µs | 140 ± 22 | 100 ± 7 | 412 ± 154 | 1323 ± 761 | |
| Percent deviation (%) | -4.5 | -25.0 | +24.1 | +6.8 | 15.1 ± 31.8 |
| Ref. *m* | 0.65 | 0.81 | 0.46 | 1.86 | |
| 150 µs< $t_d$ <400 µs | 0.62 ± 0.19 | 0.72 ± 0.17 | 0.57 ± 0.28 | 2.45 ± 1.35 | |
| Percent deviation (%) | -4.6 | -11.1 | +23.9 | +31.7 | 17.8 ± 45.9 |
| $t_d$ > 400 µs | 0.62 ± 0.12 | 0.74 ± 0.1 | 0.55 ± 0.24 | 1.75 ± 1.34 | |
| Percent deviation (%) | -4.6 | -7.4 | +19.6 | -5.9 | 9.4 ± 38.7 |

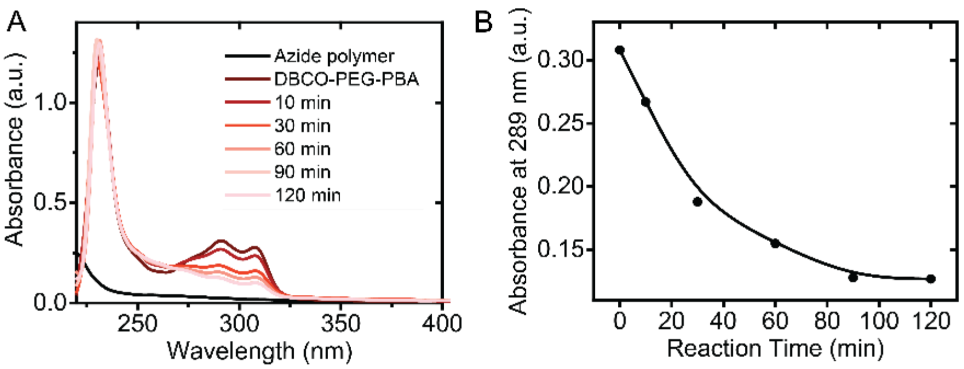


**Figure S1. UV-Vis monitoring of the reaction kinetics between PAcrAm-g-PEG-Azide and DBCO-PEG-PBA in solution. A.** UV absorption spectra of solutions containing 0.1 mg/mL PAcrAm-g-PEG-Azide only (black), 20 mg/mL DBCO-PEG-PBA only, and their mixture with 0.1 mg/mL PAcrAm-g-PEG-Azide and 20 mg/mL DBCO-PEG-PBA after increasing reaction times. **B.** Decrease of the characteristic UV absorbance peak of the DBCO group at 289 nm over time, indicating the progression of the reaction. The reaction reached a plateau after 90 min; we chose a shorter 60 min incubation time for nanopore coating to minimize the risk of pore clogging.

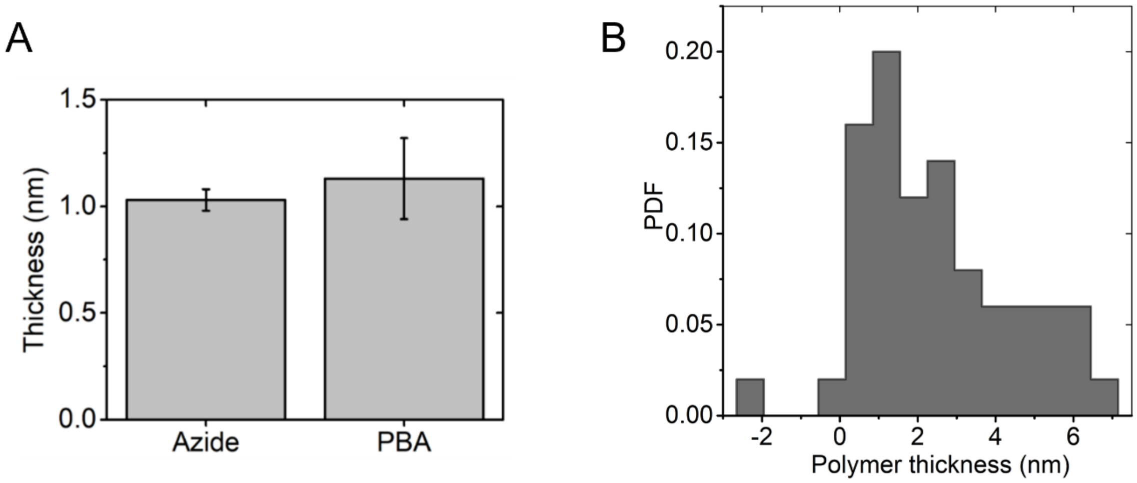


**Figure S2. Analysis of the thickness of PAcrAm-g-PEG-Azide and PAcrAm-g-PEG-PBA coatings on $SiN_x$ substrates by ellipsometry. A.** Ellipsometry measurements of a dried PAcrAm-g-PEG-PBA coating and a PAcrAm-g-PEG-Azide coating. The substrate used for coating consisted of three layers: bulk Si, 110 nm of $SiO_2$, and a 30 nm top layer of $Si_3N_4$. **B.** Histogram of the measured thickness of PAcrAm-g-PEG-PBA coating by ion conductance from nanopores of different diameters ranging from 10 to 25 nm. The average thickness and standard deviation of the PAcrAm-g-PEG-PBA coating are 1.4 ± 1.3 nm ($N$ = 50).

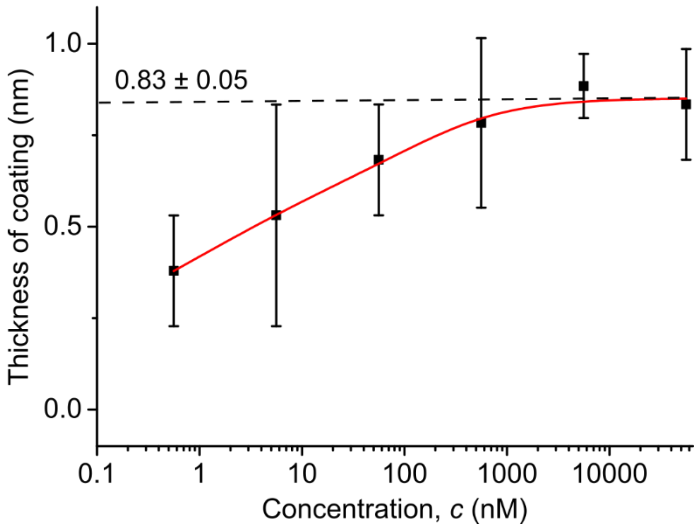


**Figure S3. Effective thickness of the glucose layer as a function of glucose concentration in a PBA-coated nanopore.** The dashed line indicates the maximum thickness of the glucose layer (0.83 ± 0.05 nm). The red curve shows the results of fitting a two-site Langmuir isotherm (Equation S3) to the coating thickness as a function of glucose concentration. Ionic current through the nanopore was recorded at -100 mV applied voltage with 500 kHz sampling rate in 2 M KCl, 10 mM HEPES, pH 7.5, at various glucose concentrations. Error bars represent the standard deviation of three independent measurements. The fitting yielded an adjusted $R^2$ of 0.99, maximum adsorption capacities of 0.32 and 0.53 nm, and dissociation constants of 71.49 and 0.235 nM.

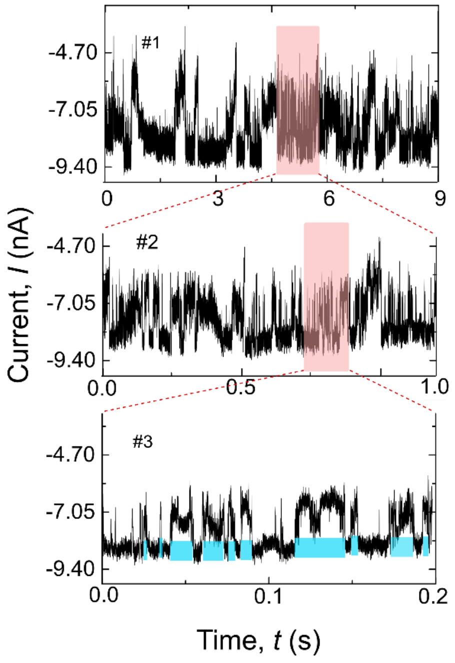


**Figure S4. Sequential zoom-in views of a representative gHSA current trace shown in Figure 2E** highlighting finer temporal features at increasing resolution. Areas shaded in red indicate the zoomed-in sections.

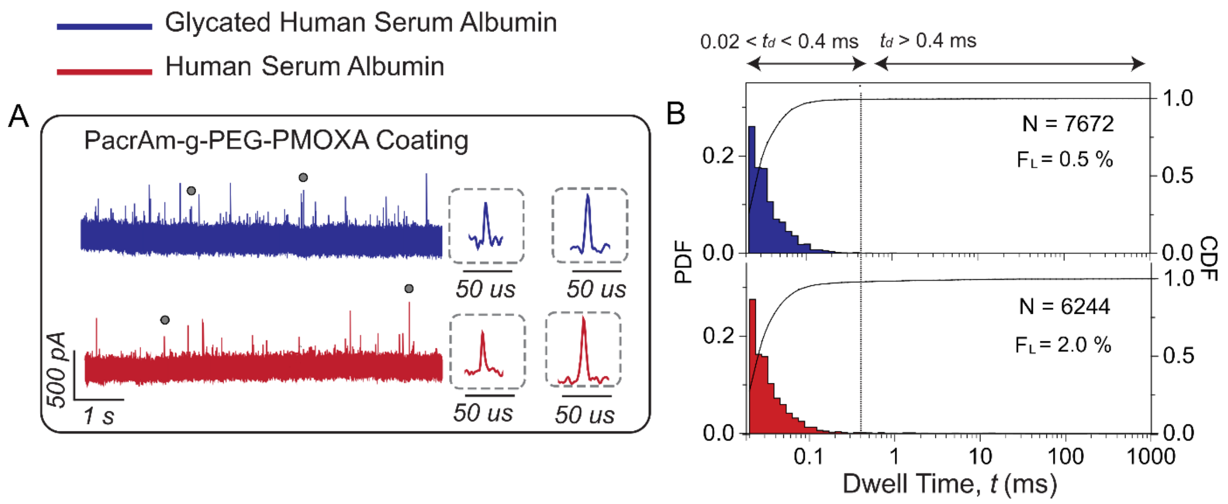


**Figure S5. Control experiment showing recordings of translocations of gHSA and HSA through PAcrAm-g-PEG-PMOXA-coated SiNx nanopore; these protein-resistant coating did not present a PBA group. A)** Representative ionic current recordings of HSA (red) and gHSA (blue) in PMOXA-coated nanopores. Insets display individual resistive pulses **B)** Probability density functions (PDF) and cumulative distribution functions (CDF) of the logarithm of dwell times from the translocation of gHSA (blue) or HSA (red). Dotted line represents the dwell time threshold of 400 µs for determining the long trapping events and free translocation events. All events that have a dwell time of at least 20 µs and are filtered with a cutoff frequency of 50 kHz with a digital Gaussian low-pass filter. This experiment reveals that coatings without PBA groups do not trap glycated (or unglycated) proteins as expected.

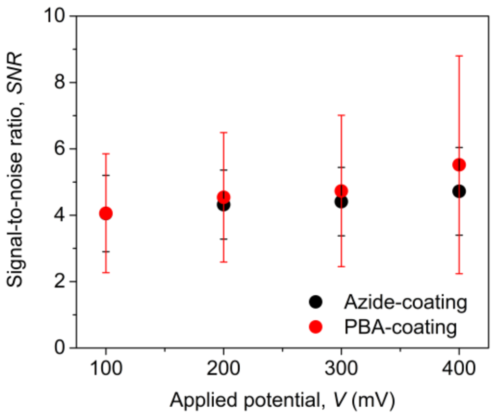


**Figure S6. Signal-to-noise ratio of ionic current obtained in the presence of gHSA as a function of the applied potential difference.** The signal-to-noise ratio for each individual resistive pulse was calculated as the ratio of the average pulse amplitude to the standard deviation of the local baseline. Only resistive pulses with dwell time of at least 20 µs were included in the analysis.

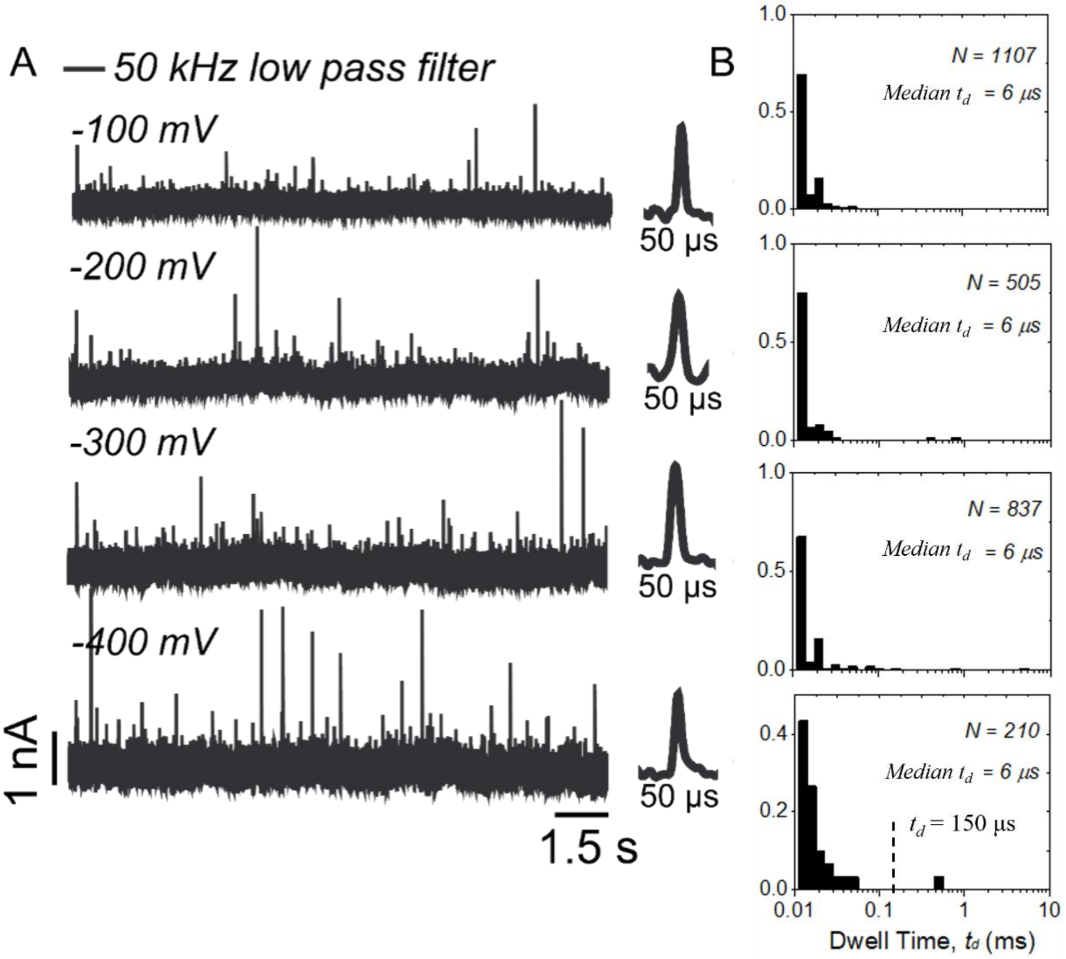


**Figure S7. Control experiment: Analysis of dwell times of gHSA in PAcrAm-g-PEG-Azide-coated nanopores at different applied voltages. A.** Representative current traces at different applied voltages, Gaussian low-pass filtered at 50 kHz. **B.** Corresponding PDFs of dwell times at different applied voltages. The median dwell time, $t_d$, was calculated without applying any dwell time thresholds. The dashed line at $t_d$ = 150 µs indicates the minimum dwell time required for volume and shape analysis. All measurements were performed with the same nanopore as in **Figure 4.**

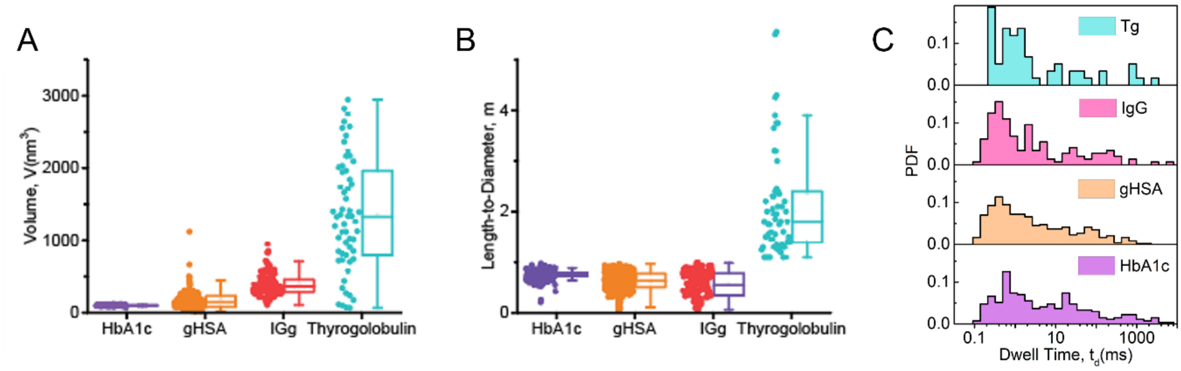


**Figure S8. Analysis of the volume and shape of four individual natively folded proteins based on resistive pulses in PBA-coated nanopores. (A)** Estimated excluded volume and **(B)** length-to-diameter ratio of four test proteins. **C.** Probability density function (PDF) of dwell times of those four test proteins. Data were collected using nanopores of different diameters as explained in the main text, at a 500 kHz sampling rate, and filtered with a 50 kHz Gaussian low-pass filter.

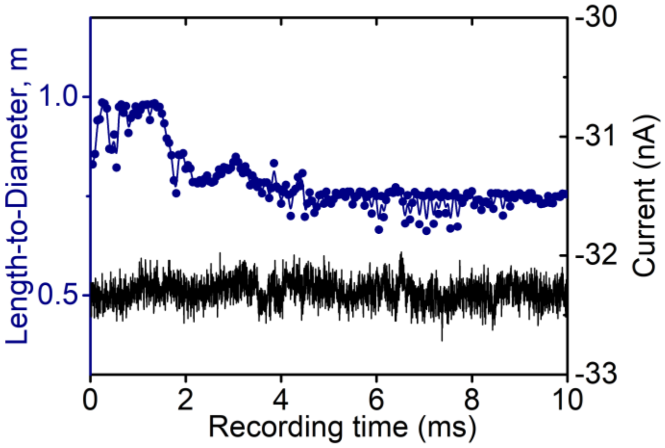


**Figure S9. The precision of the estimation of length-to-diameter ratio estimation based on a single event improves with increasing residence time of the protein in the nanopore.** The black curve is a representative current trace obtained with a PBA-coated nanopore in the presence of gHSA and Gaussian low-pass filtered at 50 kHz. The blue curve displays the results of length-to-diameter ratio estimation based on this trace as a function of increasing residence time $t_d$ in the nanopore in the range from 0 to 10 ms.

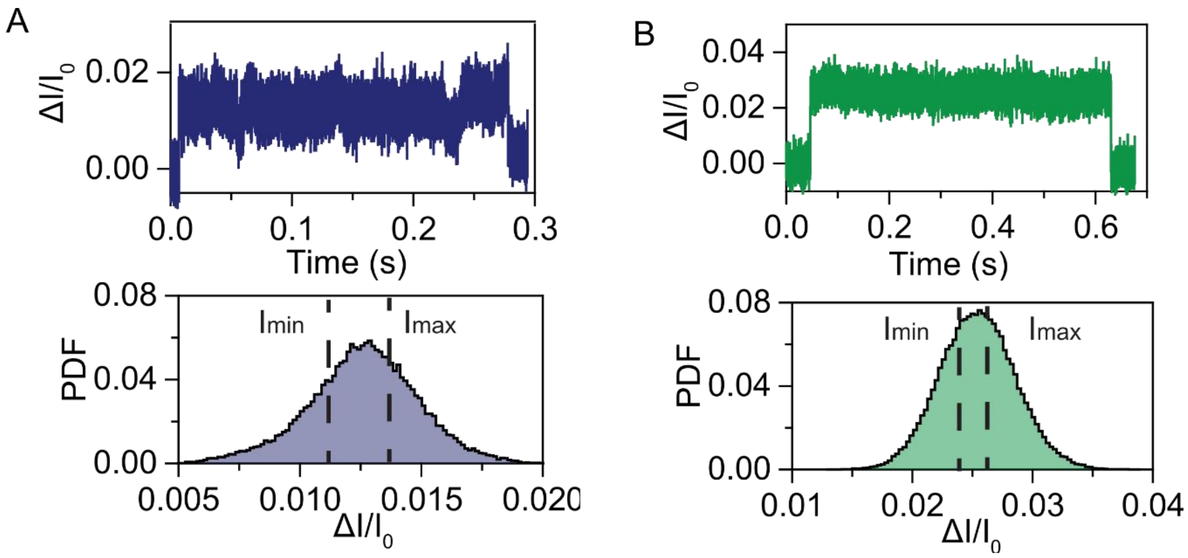


**Figure S10. Shape and volume estimation of two glycated proteins, gHSA and HbA1c, with PBA-coated nanopores. A, B:** Examples of original current traces of long trapping events and corresponding histograms of relative blockade current, $\Delta I/I_0$, for (A) gHSA, and (B) HbA1c. The $I_{min}$ and $I_{max}$ were determined by fitting the probability density distribution of the relative blockade current with **Equation S8**. The experiments were performed at -200 mV applied voltage, 500 kHz sampling rate, 50 kHz Gaussian low-pass filter in 2 M KCl, 10 mM HEPES, pH 7.4 recording buffer.
